# A Poisson reduced-rank regression model for association mapping in sequencing data

**DOI:** 10.1101/2022.05.31.494236

**Authors:** Tiana Fitzgerald, Andrew Jones, Barbara E. Engelhardt

## Abstract

Single-cell RNA sequencing (scRNA-seq) technologies allow for the study of gene expression in individual cells. Often, it is of interest to understand how transcriptional activity is associated with cell-specific covariates, such as cell type, genotype, or measures of cell health. Traditional approaches for this type of association mapping assume independence between the outcome variables (or genes), and perform a separate regression for each. However, these methods are computationally costly and ignore the substantial correlation structure of gene expression. Furthermore, count-based scRNA-seq data pose challenges for traditional models based on Gaussian assumptions. We aim to resolve these issues by developing a reduced-rank regression model that identifies low-dimensional linear associations between a large number of cell-specific covariates and high-dimensional gene expression readouts. Our probabilistic model uses a Poisson likelihood in order to account for the unique structure of scRNA-seq counts. We demonstrate the performance of our model using simulations, and we apply our model to a scRNA-seq dataset, a spatial gene expression dataset, and a bulk RNA-seq dataset to show its behavior in three distinct analyses. We show that our statistical modeling approach, which is based on reduced-rank regression, captures associations between gene expression and cell- and sample-specific covariates by leveraging low-dimensional representations of transcriptional states.

## 1 Background

Recent advances in high-throughput genomic assays have allowed for the creation of expansive data sets that are useful for exploring biological variation across cells. In particular, single-cell RNA-sequencing (scRNA-seq) technologies provide gene expression measurements at the individual cell level, allowing for the analysis of variation in transcriptional activity across cells within a single sample (Tang et al., 2009; Sasagawa et al., 2013; Jaitin et al., 2014; Zeisel et al., 2015). While characterizing this variation is useful by itself for exploratory analysis, it is also of interest to study in a more targeted way how variation relates to cell-specific covariates, such as cell type, genotype, and cell health. Studying associations between gene expression and properties of single cells has the potential to enrich our understanding of the relationship between these covariates and transcription at single-cell resolution.

While methods for association studies have been widely developed for bulk RNA sequencing (RNA-seq) data (McCarthy et al., 2008; Purcell et al., 2007), methods for studying associations on the level of individual cells are much less developed. Moreover, there are several unique challenges in manipulating and analyzing the data generated by these single-cell assays, as compared to bulk RNA-seq assays. These scRNA-seq data sets are high dimensional — there are tens of thousands of genes in the human genome — which makes them difficult to interpret gene-by-gene; furthermore, the count-based nature of the data — made up of counts of sequenced RNA fragments that map to a specific gene in a genome to approximate expression levels of that gene — presents a challenge for many standard statistical tools that make Gaussian assumptions (Cantor et al., 2010).

In this paper, we propose a statistical modeling approach based on reduced-rank regression that captures associations between gene expression and cell- and sample-specific covariates by leveraging low-dimensional representations of transcription. Within this framework, we propose two specific models: Poisson reduced-rank regression (PRRR), which adapts a generalized linear model to the reduced rank setting, and nonnegative Poisson reduced-rank regression (nn-PRRR), which provides interpretable nonnegative regression components. In what follows, we first review several related threads of research, and describe our modeling approach. Then, using simulated data and single-cell RNA-seq, bulk RNA-seq, and spatial gene expression experiments, we show that our models are useful for a wide range of association study types, including studying the transcriptional hallmarks of cell types, genotypes and eQTLs, and genes correlated with disease status.

### 1.1 Genome-Wide Association Studies

Since the completion of the Human Genome Project in 2003 (Collins et al., 2003) and the HapMap project in 2005 (Consortium et al., 2003), researchers have developed the genomic and statistical tools necessary to study the human genome at a large scale in order to better detect, treat, and prevent diseases. Genome-wide association studies (GWAS) are used to identify disease-causing genetic variation across complete genomes. Genetic variation often comes in the form of single nucleotide polymorphisms (SNPs) that can be compared between healthy patient and patients with a disease (Bush and Moore, 2012). GWAS approaches have found a plethora of gene variants that are associated with common diseases such as asthma, type 2 diabetes, and more (Ober and Nicolae, 2011; Frayling, 2007).

In a similar vein, quantitative trait loci (QTL) studies identify associations between genetic variants and quantitative phenotypes by examining molecular markers (Zeng, 1994; Doerge, 2002). A common experimental setup is to use gene expression levels as the molecular marker, in which case the study is referred to as an expression QTL (eQTL) (Nica and Dermitzakis, 2013; Kendziorski et al., 2006). Most eQTL studies have relied on bulk RNA-seq technologies to measure the gene expression levels of entire tissues (Pickrell et al., 2010; GTEx Consortium, 2017, 2020).

In this context, the statistical eQTL problem is to estimate the pairwise association between a set of genetic variants (the covariates or explanatory variables) and the expression level of each gene (the outcome variables). This is typically performed using a linear regression model. In particular, let **X** be an *N* × *P* matrix containing information about genetic variants across *P* SNPs for *N* individuals or tissue samples, and let **Y** ∈ ℝ^*N* ×*Q*^ be a matrix of corresponding gene expression levels across *Q* genes in these individuals or tissues. Eqtl approaches typically use a linear model to find associations between genotype and phenotype, estimating these relationships (Cantor et al., 2010). Generically, these approaches use a model of the form

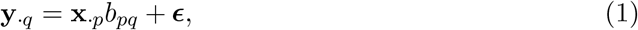

where **x**_*·p*_ is the *p*th column of **X, y**_*·q*_ is the *q*th column of **Y, *ϵ*** ∈ ℝ^*N*^ is a vector of independent zero-mean Gaussian-distributed noise terms, and *b*_*pq*_ ∈ ℝ is a scalar coefficient representing the linear relationship between SNP *p* and gene *q* for *p* = 1, …, *P* and *q* = 1, …, *Q*. Downstream tests for significance can be performed on these coefficients to identify associations (Li and Leal, 2008; Wu et al., 2011; Lee et al., 2012). Without further assumptions, this model estimates the marginal association between single SNPs and single genes independently. To accommodate polygenic contributions to phenotypes, multivariate models of the form

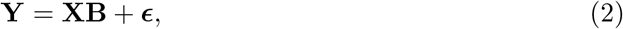

have been considered, where **B** ∈ ℝ^*P* ×*Q*^ is a matrix of coefficients (Hoggart et al., 2008; Logsdon et al., 2010; Wu et al., 2009). Under this framework, sparsity-inducing priors for **B** have been proposed in order to scale these models to high-dimensional data (Li et al., 2011, 2015).

The advent of scRNA-seq technologies has opened the door for narrowing the investigation of genotype-phenotype relationships from the level of whole tissues to the level of individual cells. However, existing computational tools are insufficient for this purpose: they typically do not accommodate count-based data, and they are seldom robust to high-dimensional outcome variables. It is difficult to control the hypothesis testing error rate in many eQTL analyses, which run millions to trillions of univariate association hypothesis tests (one for each SNP-gene pair) (Cantor et al., 2010; GTEx Consortium, 2020, 2017; Karczewski et al., 2021; Willer et al., 2013).

### 1.2 Count-based models

A further drawback of existing association testing frameworks is their assumption of Gaussian-distributed data. Most canonical regression models assume an independent normally-distributed response variable, with ***ϵ*** ∼ 𝒩 (**0**, *σ*^2^**I**_*N*_) in Equation 1. However, when the data consist of count-based measurements, this assumption may be problematic. Various transformations have been proposed to make the response variable approximately Gaussian (Butler et al., 2018a; Yu, 2009), but these transformations are known to distort the data distribution in undesirable ways (Townes et al., 2019; Booeshaghi and Pachter, 2021; Hafemeister and Satija, 2019). Count-based scRNA-seq data is discrete and nonnegative, with many gene expression counts having a value of zero. The sparsity of the data poses a challenge to these standard transformations (Townes et al., 2019).

An alternative to this approach is to model the gene expression data with a discrete distribution. A common choice is the Poisson distribution, whose support is restricted to the nonnegative integers and has been shown to improve the representation and interpretation of scRNA-seq data when fitting statistical models (Townes et al., 2019; Jones et al., 2021). A recent approach using a Poisson data likelihood proposed a naive Bayes model that assigns cell-type identities to samples in scRNA-seq data based on reference data (Grabski and Irizarry, 2020). The model uses a Poisson distribution to represent the count-based data, but the high number of zeros in the data still poses a challenge. The sparsity of the data interferes with standard estimates such as maximum likelihood estimates as rates of zero can be produced for thousands of genes, making the model sensitive to genes with low expression counts (Grabski and Irizarry, 2020). The model handles this challenge by introducing a hierarchical structure, placing posterior distributions on parameters in order to recover nonzero rate estimates for genes with zero counts in the reference data. However, the naive Bayes model also assumes independence between genes, but this assumption does not hold in practice, as expression has been observed to be correlated between genes (Butler et al., 2018b; Townes et al., 2019; Van Dam et al., 2018).

### 1.3 Modeling multiple data modalities

Latent variable modeling approaches have also been proposed for modeling multi-view data. The most popular approach has been canonical correlation analysis (CCA, (Hotelling, 1992)) and its probabilistic variants (Bach and Jordan, 2005; Zhao et al., 2016; Argelaguet et al., 2018). CCA seeks a low-dimensional linear mapping of each dataset such that the resulting low-dimensional vectors are maximally correlated.

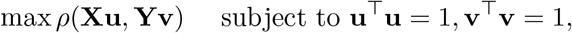

where *ρ* is the Pearson correlation function. The probabilistic version of this model projects the features of each data modality into a shared low-dimensional latent space, assuming heteroskedastic residual errors, maximizing the amount of variance explained in the data modalities by the latent subspace. The weights, or factor loadings, in CCA models allow us to identify covarying features across data modalities. A formal connection between CCA and reduced-rank regression has been shown (Tso, 1981), where the canonical subspace found by CCA is the same as the subspace of the maximum likelihood estimator for the reduced-rank regression model. Despite their connections, the unsupervised nature of CCA does not lend itself directly to association mapping between the data modalities. Conversely, reduced-rank regression has a natural association testing framework because of its regression foundation.

Recently, a latent variable model based on latent Dirichlet allocation (Blei et al., 2003; Pritchard et al., 2000) for jointly modeling gene expression and genotype was proposed (Gewirtz et al., 2021). This model projected both genotype data — using an equivalent of the Structure model (Pritchard et al., 2000) — and count-based gene expression data — using a telescoping LDA model (Blei et al., 2003) — onto a shared latent subspace; we may then identify covarying genes and gentoypes in a nonnegative latent representation. But discovering associations in this framework requires association testing in held-out data, which is limited by existing univariate methods and population data.

### 1.4 Reduced-Rank Regression Approaches

The transcriptional states of cells tend to exhibit strong correlation between genes (Stuart et al., 2003). Thus, it is likely that the relationship between cell covariates and transcriptional phenotypes in scRNA-seq data need not be modeled gene-by-gene. Rather, it is reasonable to assume that these associations exhibit low-dimensional structure. Furthermore, treating each gene as independent is computationally and statistically inefficient; we would like to exploit these relationships to perform fewer association tests and leverage shared variation to improve statistical power in these often small sample sizes. These ideas motivate a regression model whose coefficient matrix has low rank. Several approaches to reduced-rank regression have been developed to take advantage of this opportunity.

Consider again the linear regression model in Equation 2. Here, **B** is a *P* × *Q* matrix of regression coefficients, where *P* is the number of covariates, and *Q* is the number of genes. In most gene expression studies, *Q* (and sometimes *P*) is large, and min(*P, Q*) ≫ *n*. The core assumption of reduced-rank regression (RRR) is that the matrix **B** has low rank (Anderson et al., 1951). In particular, the RRR model assumes **B** has rank *R* ≪ min(*P, Q*). This implies that **B** can be factorized as an outer product of two low-rank matrices, giving us the following reduced-rank regression model:

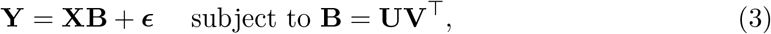

where **U** ∈ ℝ^*P* ×*R*^ and **V** ∈ ℝ^*Q*×*R*^. In the context of gene expression studies, this low-rank assumption implies that the relationship between cell-specific covariates and gene expression can be described in terms of a small set of latent factors. In other words, variance in gene expression is mediated by *R* different programs encoded in subsets of covariates; then **B** captures both the covariates of interest and their effect sizes within each of the *R* programs.

Several estimation approaches have been proposed for RRR under the assumption of Gaussian noise. A common method is to find the parameter values that minimize the squared reconstruction error (Anderson et al., 1951; Reinsel and Velu, 1998):

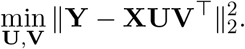

This approach corresponds to finding the maximum likelihood solution of an RRR model with Gaussian errors (***ϵ***_*q*_ ∼ 𝒩 (0, *σ*^2^**I**) for *q* = 1, …, *Q* in Equation 3 as *σ*^2^ → 0). When **B** is assumed to have full rank (that is, *R* = min(*P, Q*)) the minimization admits the ordinary least squares (OLS) solution:

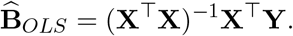

When *R <* min(*P, Q*), the RRR model has an eigenvalue solution:

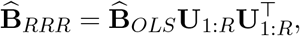

where 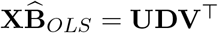 is the SVD of the fitted values, and **U**_1:*R*_ = [**u**_1_, · · ·, **u**_*R*_] contains the leading *R* left singular vectors of 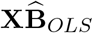.

Sparse approaches to RRR have been proposed as well. Sparsity in the decomposition leads to greater interpretability by including nonzero weights only on a subset of the covariates and genes for any component. One model (Qian et al., 2020) imposes sparsity on the coefficient matrix **B** by taking an iterative approach to estimation, solving both a sparse regression problem and the reduced-rank decomposition in alternating frames. The base algorithm solves the following optimization problem:

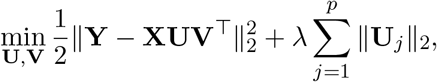

where **U** ∈ ℝ^*P* ×*R*^, **V** ∈ ℝ^*Q*×*R*^, **U**_*p*_ represents the *p*th row of **U**, the rank *R* is specified by the modeler, and *λ* is a sparsity penalty parameter. The alternating minimization problem can be broken into two steps: optimizing **U**, and optimizing **V**. After parameter initialization on iteration 𝓁 = 1, on iteration 𝓁 = 2, …, *L*, the algorithm first solves an orthogonal Procrustes problem for **V**:

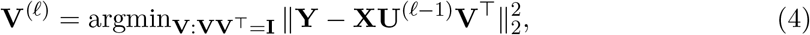

where **U**^(𝓁*−*1)^ is the estimate of **U** from the previous iteration. The algorithm then solves a group lasso problem for **U**:

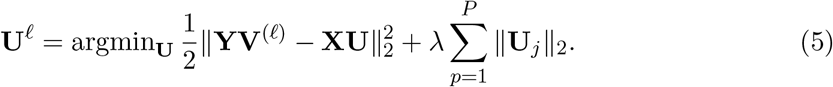

Equation 4 can be solved using a singular value decomposition, and Equation 5 can be solved using techniques for group lasso (Friedman et al., 2010). These two steps are repeated for *L* steps or until convergence.

Another approach developed a Bayesian RRR framework for association mapping in the GWAS setting (Valente et al., 2015). The model — called Bayesian Extendable Reduced-Rank Regression (BERRRI) — uses a nonparametric Indian Buffet Process prior for the latent factors, which allows the rank *k* to be estimated from the data. BERRRI then uses a variational Bayes approximation to the posterior for inference of the model parameters. However, BERRRI does not explicitly model count-based data, and its inference procedure is not computationally tractable for genome-scale analyses.

The linear RRR model has been generalized to nonlinear functions as well. The most popular nonlinear approaches have used neural networks with multiple inputs and multiple outputs (Diamantaras and Kung, 1994). The linear RRR model is equivalent to a single-layer multi-layer perceptron with only linear transformations between layers (Baldi and Hornik, 1989; Kunin et al., 2019). This model can be extended to the nonlinear case by including nonlinear activation functions (Baldi and Hornik, 1989; Aoyagi and Watanabe, 2005). However, these models typically to do not capture count data and lack the interpretability of linear models for downstream association testing.

In this manuscript, we propose a statistical model and associated computational framework that addresses the problems that arise with modeling genotype-phenotype associations for high-dimensional phenotypes captured with count data. We propose a reduced-rank regression model that finds low-dimensional associations between genotypes (or other high-dimensional covariates) and and RNA-sequencing data (or other high-dimensional count-based phenotypes). Relying on low-dimensional associations alleviates the problem of estimating millions of pairwise associations. Furthermore, our model uses count-based likelihoods that allows both single-cell RNA-sequencing data and bulk RNA-sequencing. We show that our approach appropriately models gene expression data with count-based likelihoods, leads to interpretable subsets of genes and genetic variants in each dimension, and uses flexible, computationally tractable inference methods that allow for uncertainty quantification.

## 2 Methods

We propose a probabilistic reduced-rank regression model with a Poisson data likelihood — which we call Poisson reduced-rank regression (PRRR) — for association mapping in count-based sequencing data. Our approach takes the form of a reduced-rank regression model with intermediate factors that explicitly model count-based data using a Poisson likelihood. These factors are interpretable and can be used to to identify and analyze the global structure of associations between cell covariates and cell phenotypes, such as gene expression levels. We ensure that inference is tractable and efficient in these models by leveraging a stochastic variational inference approach.

### 2.1 Poisson Reduced-Rank Regression (PRRR)

PRRR is designed to identify associations between cell-specific covariates and high-dimensional gene expression profiles. The response matrix 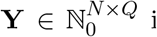 is a matrix containing (in this application) RNA transcript counts for *Q* genes in *N* cells, where ℕ_0_ = ℕ ∪ 0. The *N* × *P* matrix **X** is a design matrix containing covariates for each cell. For example, these covariates could represent cell type, genotype, or measures of cell health.

PRRR uses a Poisson likelihood to model the transcript counts for each cell as the response variables, conditional on observed cell-specific covariates. The Poisson rate is parameterized by a low-rank linear mapping from the cell covariates.

Specifically, the transcript count of gene *p* in cell *n*, denoted by *y*_*np*_ is modeled as a draw from a Poisson distribution, *y*_*np*_ ∼ Poisson(*λ*_*np*_). The Poisson rate *λ*_*np*_ is determined by a linear function of the vector of covariates for cell *n*, denoted as **x**_*n*_. We use a canonical link function from the exponential family to map the domain of the latent variables to the positive real line — similar to a GLM approach. In particular, we use a log link function to ensure that, when pushed through the inverse link — the exp function — the linear predictor lies in R_+_. The likelihood model is then

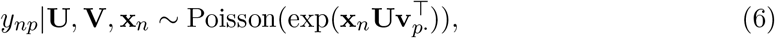

where **v**_*p·*_ is the *p*th row of **V**. We place Gaussian priors on columns of **U** and **V**:

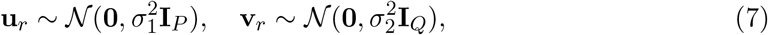

for *R* = 1, …, *R*. Intuitively, **U** and **V** capture the low-rank associations between **X** and **Y**.

#### 2.1.1 Nonnegative PRRR

In some cases, the covariates **X** are entirely nonnegative — possibly representing counts or categories — in which case it may be of interest to identify nonnegative, low-rank regression coefficients that explain the associations in a completely additive fashion. For example, in eQTL mapping, the covariates are typically the count of the minor allele for each SNP, where **x**_*n*_ ∈ {0, 1, 2}, and it may be of interest to identify a nonnegative, “parts-based” combination of SNPs that explain phenotypic variation. For these cases, we propose nonnegative Poisson reduced-rank regression (nn-PRRR), whose likelihood is given by

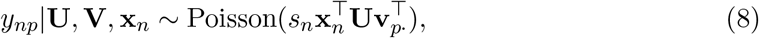

where *s*_*n*_ is a cell-specific size factor modeling the total number of transcripts in cell *i*. We fix *s*_*n*_ to be the total number of transcript counts in cell, 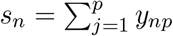. We place Gamma priors on the elements of **U** and **V**:

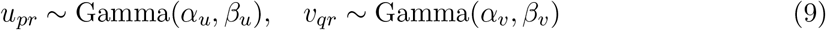

for *p* = 1, …, *P, q* = 1, …, *Q*, and *R* = 1, …, *R*. For all experiments, we set *α*_*u*_ = *α*_*v*_ = 2 and *β*_*u*_ = *β*_*v*_ = 1.

### 2.2 Estimation and inference

We propose two approaches to fit our model to data: 1) computing a point estimate for the coefficients using maximum *a posteriori* (MAP) estimation and 2) full Bayesian posterior inference for the regression coefficients using an approximate inference procedure.

#### 2.2.1 MAP estimation

The MAP solution in our model is the coefficient matrices **U**_*MAP*_, **V**_*MAP*_ with maximum posterior probability given the data **X** and **Y**. In particular,

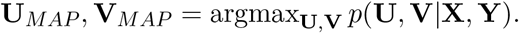

Expanding the posterior with Bayes’ rule, we can write the MAP objective as

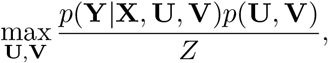

where *Z* is a normalizing constant that does not depend on **U** or **V**. Taking a log, dropping the constant *Z*, and leveraging the i.i.d. assumption, we arrive at our final MAP objective:

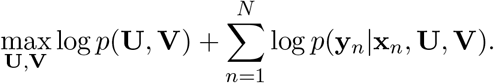

Although this maximization problem does not have a closed-form solution, we use gradient-based methods to iteratively maximize this log posterior with respect to **U** and **V**.

### 2.3 Variational inference

A fully Bayesian approach to inference, given a set of samples with paired cell covariates and transcript counts, 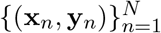, would compute the posterior distribution of the parameters, **U** and **V**, given the data matrices **X** and **Y**. By Bayes’ rule,

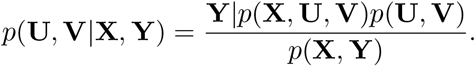

However, the marginal likelihood, *p*(**X, Y**), contains an intractable integral,

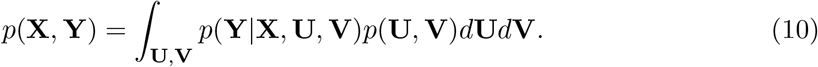

To circumvent this issue, we use a variational approximation to the posterior. Specifically, we use a mean-field variational approximation, *p*(**U, V**) ≈ *q*(**U, V**) = *q*_1_(**U**)*q*_2_(**V**), where *q*_1_ and *q*_2_ are the variational distributions. Here, we specify the variational families for PRRR and nn-PRRR to be normal and log normal, respectively,

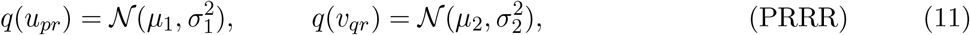

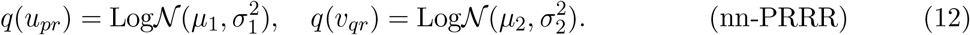

We minimize the KL divergence between the true posterior and the variational approximation, which is equivalent to maximizing a lower bound on the model evidence (called the ELBO). This lower bound for PRRR is given by

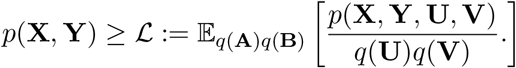

We maximize this lower bound with respect to the variational parameters using stochastic variational inference (Hoffman et al., 2013) as implemented in TensorFlow Probability (Dillon et al., 2017). For all experiments, we use the Adam optimizer (Kingma and Ba, 2014) with a learning rate of 0.01.

## 3 Results

### 3.1 Simulation experiments

We first demonstrate the use cases of PRRR and test the robustness and accuracy of our model using simulated data.

#### 3.1.1 PRRR identifies low-dimensional association maps

We first sought to confirm that PRRR identifies the low-dimensional relationships between covariates and outcomes.

To start in a setting that can be visualized, we generated a synthetic dataset in which the covariates and outcomes are both two-dimensional. Specifically, we sampled data from the generative model (Equations (8)-(9)), setting *R* = 1. We forced a correlation between the covariates and outcomes. We found that PRRR could reliably detect the one-dimensional association between **X** and **Y** (Figure 1). Moreover, we are able to recover a quantification of the relationship between the covariates and outcomes, and visualize this relationship in the low-dimensional space.

**Figure 1:**
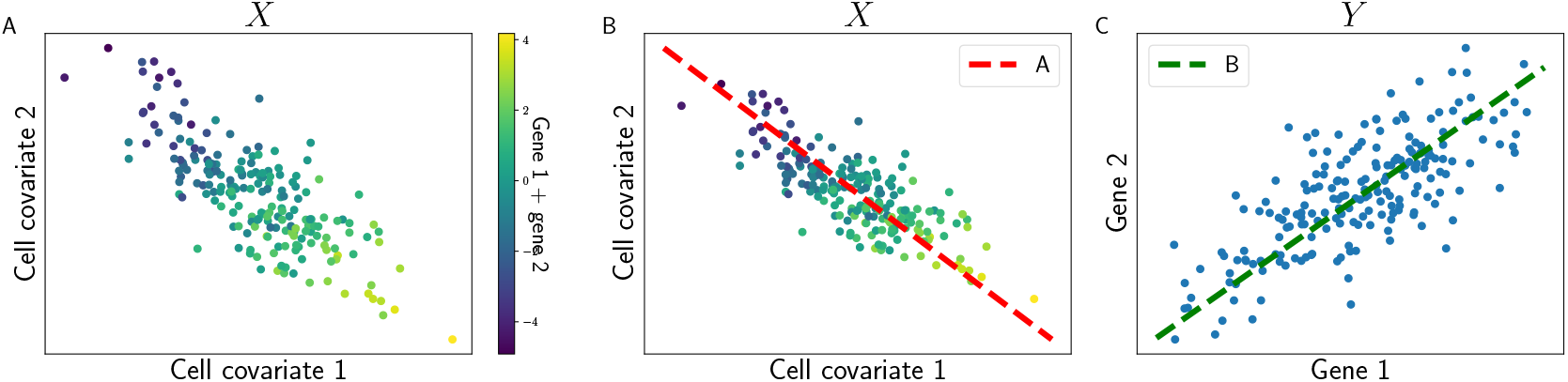
Illustration of PRRR. We fit PRRR to a toy dataset containing two cell-specific covariates and two genes. The two covariates showed negative correlation, and the two genes were jointly associated with the covariates (panel A). PRRR identifies the low-rank structure of these multivariate relationships by decomposing the full coefficient matrix into two low-rank matrices, *A* and *B* (panels B and C).

We next extended this simulation study and visualization to a small-scale synthetic eQTL study. We generated *N* = 200 synthetic genotypes based on minor allele counts, **x**_*n*_ ∈ {0, 1, 2}, and sampled synthetic RNA transcript counts using the PRRR generative model, with *P* = *Q* = 2 for visualization. We fit PRRR to these data and inspected the fitted coefficients. We found that PRRR recovered these genotype-expression relationships, and allowed for both inspection of the low-dimensional structure of these relationships, as well as investigating univariate relationships (Figure 2). This experiment suggests that PRRR may be useful to perform eQTL mapping.

**Figure 2:**
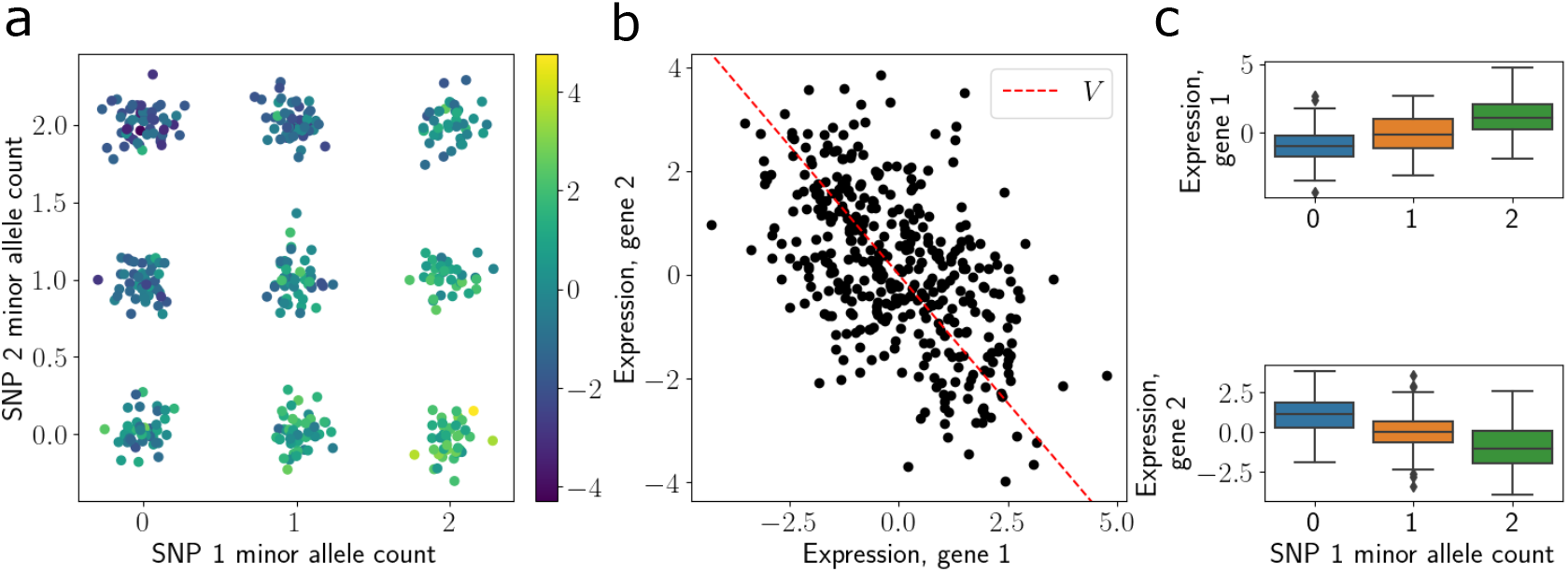
eQTL mapping in simulated single-cell data with PRRR. Toy example demonstrating eQTL mapping with PRRR for two genetic variants and two genes. (a) Genotype data, shown as the number of copies of the minor allele for variant 1 (x-axis) and variant 2 (y-axis) and colored by each sample’s corresponding expression of gene 1. (b) Gene expression values. The red line represents the fitted value for **V** with *r* = 1 in this toy example for gene 1 (x-axis) and gene 2 (y-axis). (c) Relationship between genotype (x-axis) and gene expression (y-axis) for the two genes.

#### 3.1.2 PRRR identifies the optimal rank and is robust to misspecification

PRRR, like other reduced-rank regression approaches, requires selecting the rank *R* of the coefficient matrix. A common approach is to evaluate the goodness-of-fit of the model at varying values of *R*, and select the one with the best fit to the data. To test whether this is feasible with PRRR, we use synthetic data that was generated from PRRR’s generative model with true rank *R*^⋆^ = 3. We then fit the model with *R* ∈ {1, 2, …, 10} and compute the ELBO for each fit. We repeated this experiment 30 times for each value of *R*.

We find that the PRRR ELBO peaked at the true value of *R* = 3 (Figure 3a), demonstrating that the model’s fit to the data was best at the true rank *R*^⋆^. Moreover, we found that while the goodness-of-fit sharply degraded for models with *R < R*^⋆^, the goodness-of-fit declined more slowly for models with *R > R*^⋆^. This finding confirms similar observations from previous studies (Qian et al., 2020), and suggests that setting the rank to be higher than anticipated is protective against model misspecification.

**Figure 3:**
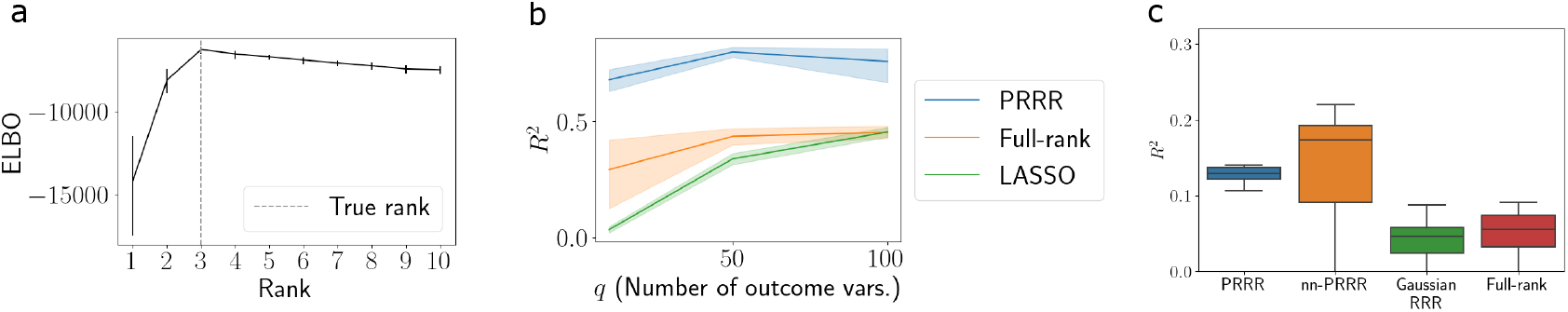
PRRR identifies optimal rank and is robust to data dimension. (a) Using synthetic data generated under the PRRR model with a true rank of *R*^⋆^ = 3, we fit PRRR with a range of rank specifications on the x-axis, where *R*^⋆^ = 3. The y-axis shows the ELBO values for each model rank. Vertical ticks represent 95% confidence intervals. (b) Goodness-of-fit *R*^2^ values for predictions from PRRR, a full-rank version of PRRR, and a multi-output LASSO model (Friedman et al., 2010) for outcome data with dimension *Q* ∈ {10, 50, 100}. (c) Goodness-of-fit *R* values for predictions from PRRR, nn-PRRR, and competing models for outcome data generated from Splatter (Zappia et al., 2017).

#### 3.1.3 PRRR is robust to data dimension

We next sought to validate the robustness of PRRR in the presence of higher-dimensional data. To do so, we generated three datasets from the PRRR model, each with a different number of response features (genes), *Q* ∈ {10, 50, 100}. We set the sample size to *N* = 200, and we randomly selected 80% of these samples to fit the model and held out the remaining 20% to test the model. We fit PRRR on the training data using our MAP estimation procedure and used the estimated parameters to compute the predicted Poisson rate for the held-out data, 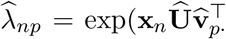. We then computed the goodness-of-fit *R*^2^ measure between our predictions and the held-out dataset’s counts. To benchmark these predictions, we compared PRRR’s predictive performance to two competing methods: a full-rank version of PRRR and a multi-output LASSO model, as implemented in the R package glmnet (Friedman et al., 2010). We repeated this experiment ten times for each method and each data-generating condition.

We found that PRRR reliably achieves good predictive performance across all values of the outcome dimension *Q* (Figure 3b). In contrast, the full-rank and LASSO models performed worse.

To further validate PRRR and nn-PRRR on different data types, we conducted a similar experiment with synthetic data generated using Splatter (Zappia et al., 2017), a data simulator designed specifically for single-cell count data. We generated data for *N* = 200 samples, each belong to one of 10 groups, and used *Q* = 100 genes. We used the one-hot encoded group labels as the covariates and the synthetic gene expression as the response. We fit PRRR and nn-PRRR with *R* = 5, along with Gaussian RRR and full-rank Poisson regression, and we reserved a hold-out set for evaluating predictions. We found that PRRR and nn-PRRR outperformed competing models in terms of their predictive *R*^2^ values (Figure 3c).

These results suggest that accounting for the count-based data and the low-rank structure of associations is vital, and that the PRRR model successfully captures this structure.

#### 3.1.4 PRRR predictions are robust to rank misspecification

To further explore the role of rank specification in our model, we performed a prediction experiment for varying settings of the rank. We generated synthetic data as before with *R*^⋆^ = 3 and fit the model on 80% of the data while reserving 20% for testing. For a range of ranks, *R* ∈ {1, 2, 3, 4, 5, 10, 20}, we fit PRRR, used the fitted model to make predictions for the held-out data, and computed the *R*^2^ coefficient of determination for these predictions. We performed this experiment using both maximum a posteriori (MAP) estimation and variational inference to fit the model, repeating the experiment ten times for each rank in both cases.

We found that the predictive performance was strongest when the model was correctly specified (*R* = 3 in this case; Figure 4a, b). However, we observed that performance was strong across a range of misspecified ranks as well. Similar to our previous experiment, we observed that predictions were more robust for models with *R > R*^⋆^ as compared to models with *R < R*^⋆^.

**Figure 4:**
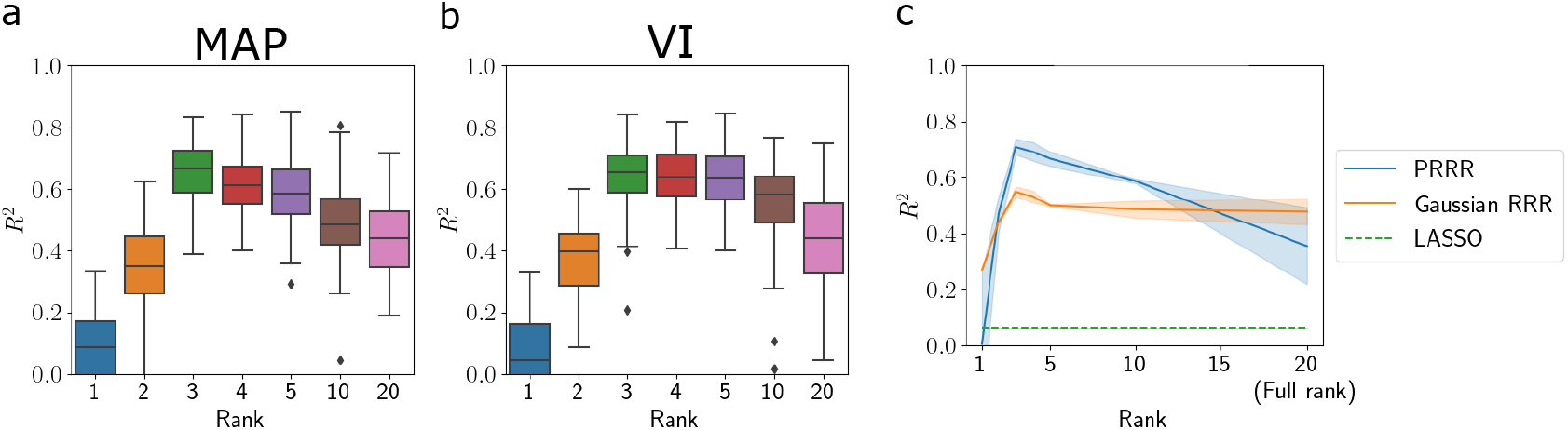
PRRR is robust to rank misspecification. Using synthetic data generated from the PRRR model with a true rank of *R*^⋆^ = 3, we fit PRRR with a range of rank specifications (*x*-axis). We made predictions for a held-out dataset and computed the *R*^2^ coefficient of determination, repeating this ten times for each rank. The *y*-axis shows the *R*^2^ value between the predicted values and the true values on held-out samples. Boxes show the median and upper and lower quartiles, and whiskers extend to 1.5 times the interquartile range. (a) Maximum *a posteriori* estimates (MAP); (b) Variational inference (VI); (c) Comparison with Gaussian RRR (Anderson et al., 1951) and LASSO (Friedman et al., 2010).

To benchmark these predictions, we compared PRRR’s predictive performance to two competing methods: a reduced-rank regression that assumes a Gaussian likelihood (Anderson et al., 1951) and the multi-output LASSO model (Friedman et al., 2010). We performed a similar prediction experiment as above, computing the *R*^2^ for each model under a range of rank specifications. We found that PRRR outperformed the two competing methods in a range around the true rank *R*^⋆^ = 3 (Figure 4c).

### 3.2 Characterizing transcriptional hallmarks of pancreatic cell types

It has been widely observed that cell type is a major driver of transcriptional variation between cells in a variety of tissue types (Zheng et al., 2017; Kotliar et al., 2019; Zeisel et al., 2015; Chen et al., 2017). Given these observed differences between cell types, it is of interest to identify the gene expression patterns that are characteristic of each cell type. Our PRRR models present a principled approach for identifying these transcriptional hallmarks of cell types by finding a low-dimensional mapping from cell type to expression.

To test this, we fit PRRR to an scRNA-seq dataset containing 1,578 cells that span *C* = 14 unique cell types in the human pancreas (Baron et al., 2016). For cell *n*, we encode its cell type as a one-hot vector **x**_*n*_ ∈ {0, 1}^*C*^, and we treat the response variable **y**_*n*_ as the vector of RNA transcript counts in this cell. We extracted the coefficient matrices **U** and **V** and studied their properties.

We found that PRRR was able to identify transcriptional markers in each cell type. Among the 14 unique cell types present in the dataset, there are five that belong to the family of islet cells (*alpha, beta, gamma, delta*, and *epsilon* cells). Given their functional relatedness, these cell types are expected to show similar gene expression patterns compared to patterns found in other cell types. Indeed, inspecting PRRR’s estimated coefficients, we find that the model captures the low-dimensional gene expression patterns in islet cells (Figure 5). We performed a hierarchical clustering on the PRRR coefficients, which revealed that the islet cells clustered together (Figure 5). We found a similar clustering after fitting nn-PRRR on the same dataset (Supplementary Figure 12). Moreover, we observed that the models separated islet cell types from non-islet cell types in the low-dimensional space (Figure 6). Examining the factor loadings for each cell type, we found that specific factors were especially enriched for islet or non-islet cell types (Figure 7). The islet-related factors were enriched for pancreatic gene pathways, such as *pancreas beta cells*.

**Figure 5:**
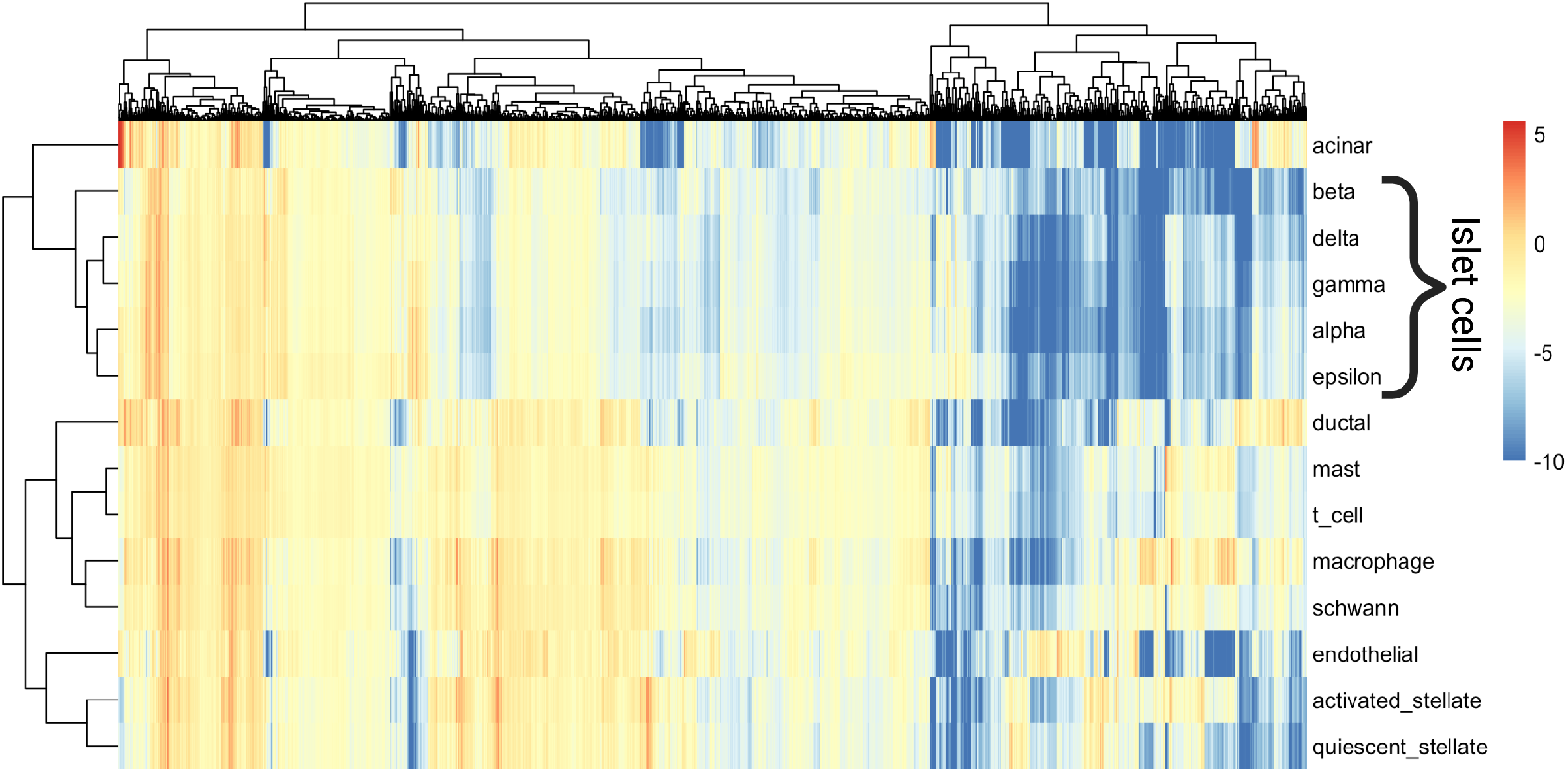
PRRR coefficients for pancreatic cell types. Heatmap showing the full coefficient matrix **UV**^*T*^, with cell types on the rows and genes on the columns.

**Figure 6:**
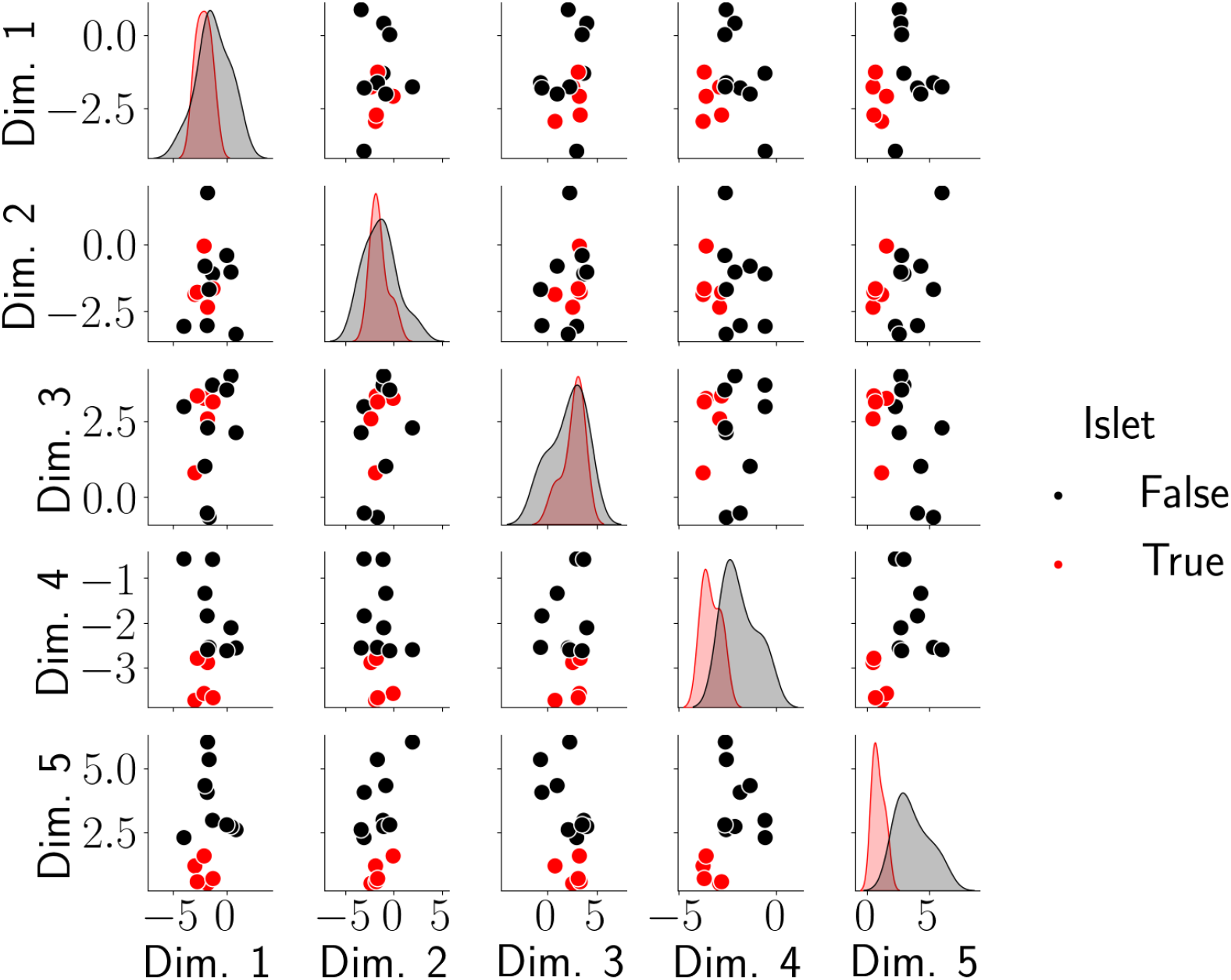
PRRR identifies similar expression patterns in islet cell types. Shown here is the latent encoding of each cell type for each pair of latent variables in **U**, where PRRR was fitted with *R* = 5 Each point in each subplot represents a cell type, and cell types are colored by whether they are classified as islet cells or not. The densities on the diagonal show the distribution of **U** values for islet and non-islet cell types in each latent dimension.

**Figure 7:**
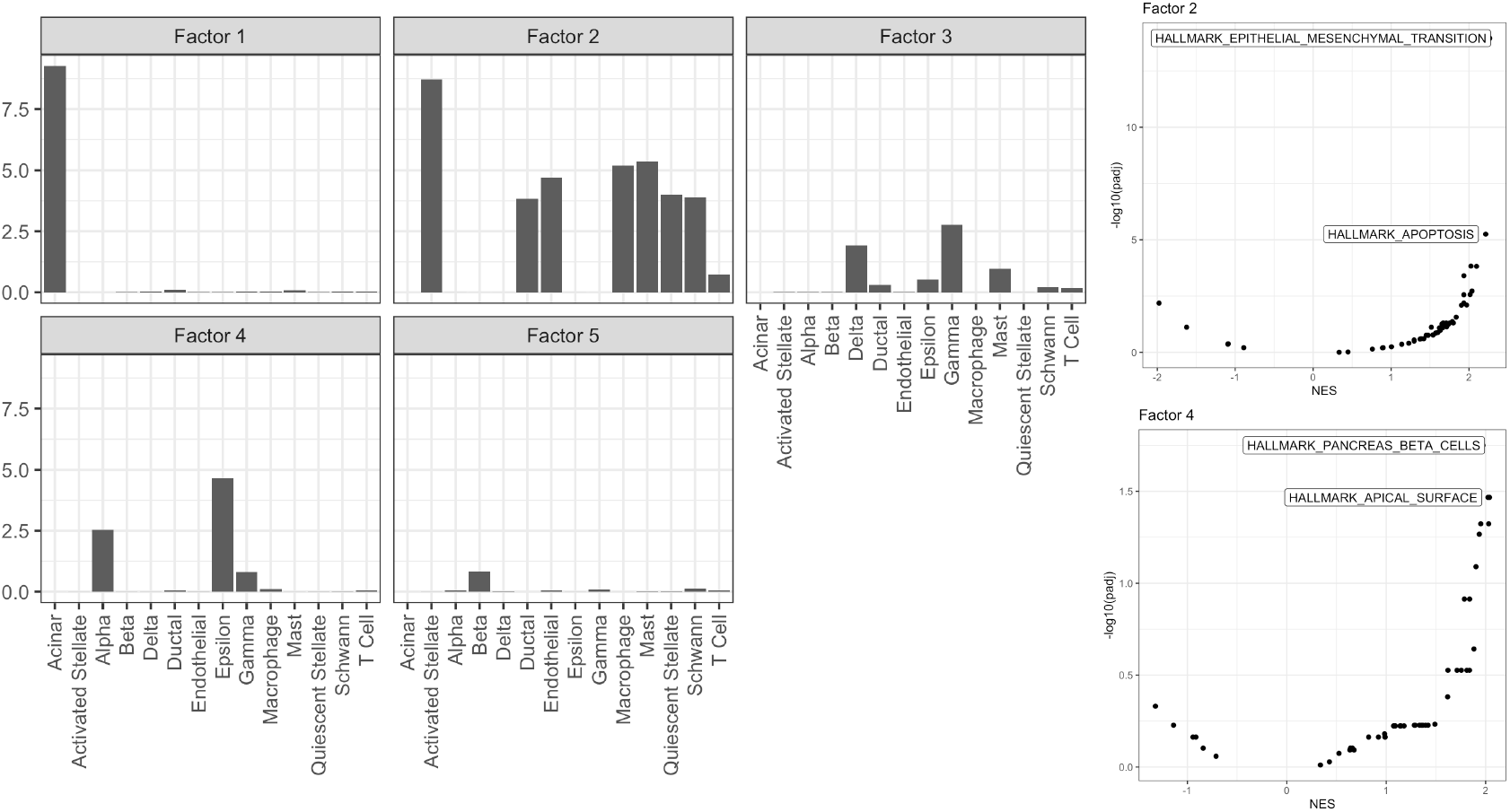
PRRR factors identify subgroups of cell types. (a) shows each cell type’s loading onto each of the five latent factors in the **U** matrix. (b) shows a gene set enrichment analysis of the gene loadings onto factors 2 and 4 in the **V** matrix.

We also extracted the top genes for each cell type from the full PRRR coefficient matrix; these genes can be viewed as “marker genes” whose expression is correlated with certain cell type identities. We found that these marker genes corresponded to cell-type-specific transcription patterns, such as *REG1B* being the top marker gene for *acinar* cells (Figure 13).

These findings imply that the low-dimensional space may be used to compare and contrast existing classifications within the data, as well as to discover possible new relationships among covariates and phenotypes. They also show that PRRR is able to identify the distinct transcriptional characteristics of specific cell types and groups of cell types, and that PRRR could be used to identify marker genes for cell types.

### 3.3 Analyzing gene expression patterns in spatial datasets

Along with cell type, the physical organization of cells within a tissue has a strong impact on gene expression due to tissue organization structure and cell-cell communication. The rise of spatial gene expression profiling technologies provides an opportunity to study how gene expression levels vary spatially across a tissue (Stickels et al., 2021; Ståhl et al., 2016; Rodriques et al., 2019; Lee et al., 2021). In particular, given gene expression data at the individual cell level with appropriate spatial context, it is of interest to identify how the expression of specific genes varies across the expanse of a tissue.

To study this with our modeling framework, we fit PRRR with rank *R* = 1 to a two-dimensional spatial dataset containing 2,063 mouse brain sagittal anterior cells (10x Genomics, 2020). The **X** matrix is an *N* × 2 matrix containing two-dimensional spatial coordinates for each cell *n*, and we treat the response variable **Y** as the matrix of RNA transcript counts. After fitting PRRR, we extracted the model coefficients to inspect the spatial trends in gene expression that it identified.

We found that PRRR model is able to identify trends in gene expression along one latent dimension. While the model is constrained to only identify linear changes in gene expression across space, it is able to identify a general trend in increased gene expression for individual genes such as *TTR* and *FABP7* (Figure 8). This result demonstrates the utility of our model in the context of spatial genomics and further demonstrates the versatility of PRRR.

**Figure 8:**
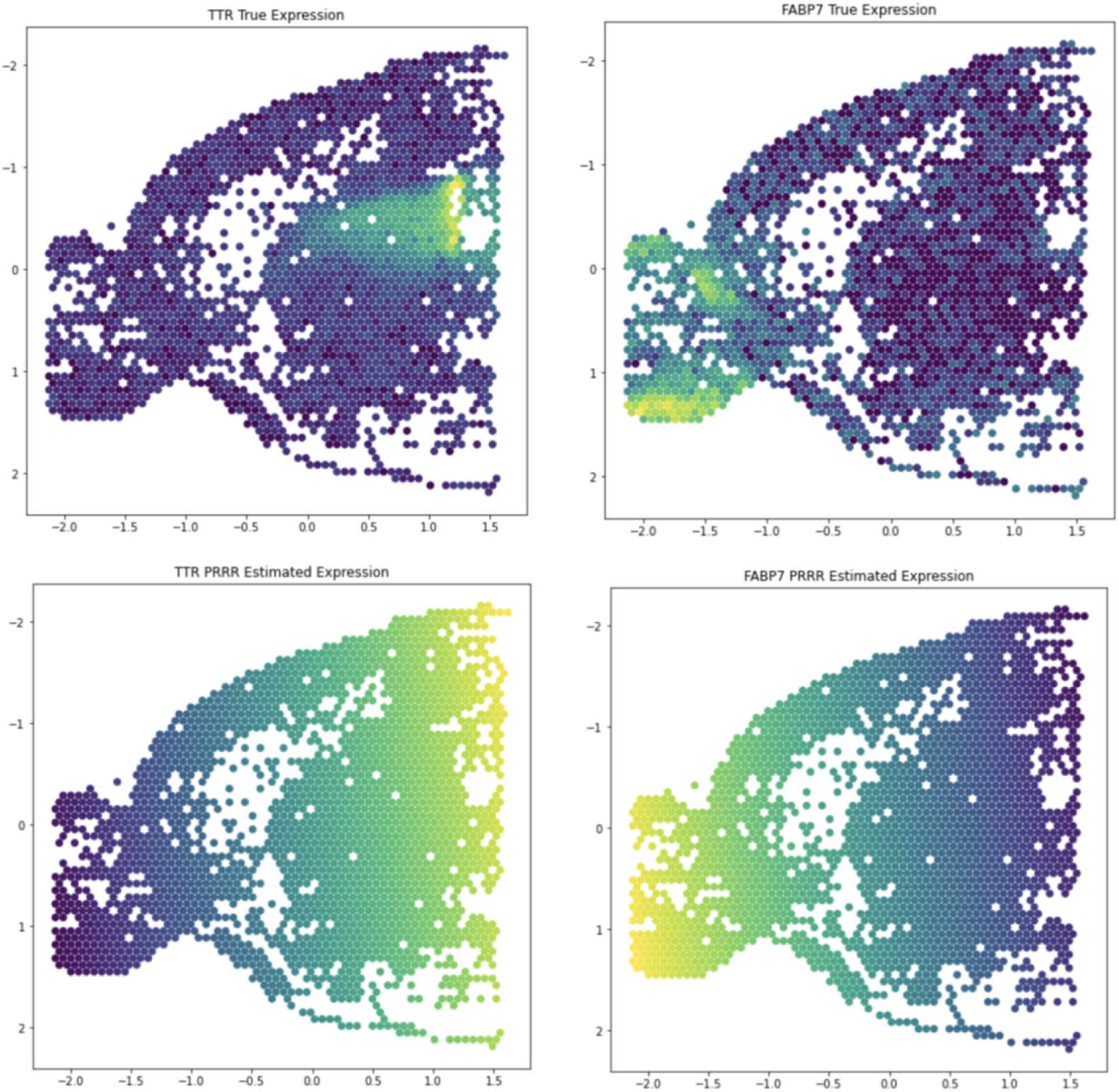
PRRR identifies directional patterns in spatial gene expression data. We applied PRRR to a Visium spatial gene expression readout from the sagittal anterior region of a mouse brain, using each spot’s spatial coordinates as the covariates and each spot’s gene expression levels as the outcome. Left: Spatial gene expression data for the gene *TTR* and PRRR’s estimated spatial pattern for this gene. Right: Spatial gene expression data for the gene *FABP7* and PRRR’s estimated spatial pattern for this gene.

### 3.4 eQTL mapping

eQTL mapping is a common approach to finding associations between a genotypes and gene expression profiles. However, this type of association mapping requires fitting a regression between millions of genotype variants and the expression of tens of thousands of genes, resulting in billions of univariate models (GTEx Consortium, 2017). The large number of univariate tests can cause these approaches to be prohibitive computationally and to lack sufficient statistical power.

We hypothesized that our reduced-rank regression model could alleviate these issues of computational tractability and statistical power. To test this, we applied PRRR to an eQTL mapping setting to find a set of low-dimensional factors that capture the relationships between genotype and gene expression. To do so, we used data from the Genotype Tissue Expression (GTEx) Consortium (GTEx Consortium, 2020). For this experiment, we focused on data collected from liver tissues from 227 donors. For each donor, the data consist of paired genotype (as encoded by minor allele count **x**_*n*_ ∈ {0, 1, 2}) and bulk gene expression profiles.

We fit PRRR with *R* = 10 latent dimensions and extracted the low-rank regression coefficient matrix (Figure 9a, b). Within each factor, we can examine associations between individual genetic variants and the expression of individual genes. For factor number *r*, we do this by taking the outer product of the corresponding columns of **U** and **V**, respectively. That is, we compute 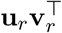 and examine the strongest SNP-gene relationships (Figure 10). Investigating the latent factors more closely, we found several meaningful associations. For example, after performing a gene set enrichment analysis, we found that the gene expression loadings for factor 9 were enriched for genes related to *interferon gamma response* and *inflammatory response* (Figure 9c), two major functional roles of liver cells (Horras et al., 2011; Robinson et al., 2016). We find similar results for nn-PRRR, although the **V** matrix is much more sparse (Figure 15). The additional sparsity in the nn-PRRR results aligns well with the parts-based representation of the low-dimensional space known with nonnegative matrix factorizations (Lee and Seung, 1999; Donoho and Stodden, 2003; Townes and Engelhardt, 2021). This experiment suggests that the PRRR and nn-RRR framework may be useful for studying associations between genotypes and phenotypes, especially when there is low-dimensional correlation structure between the datasets and within each dataset individually.

**Figure 9:**
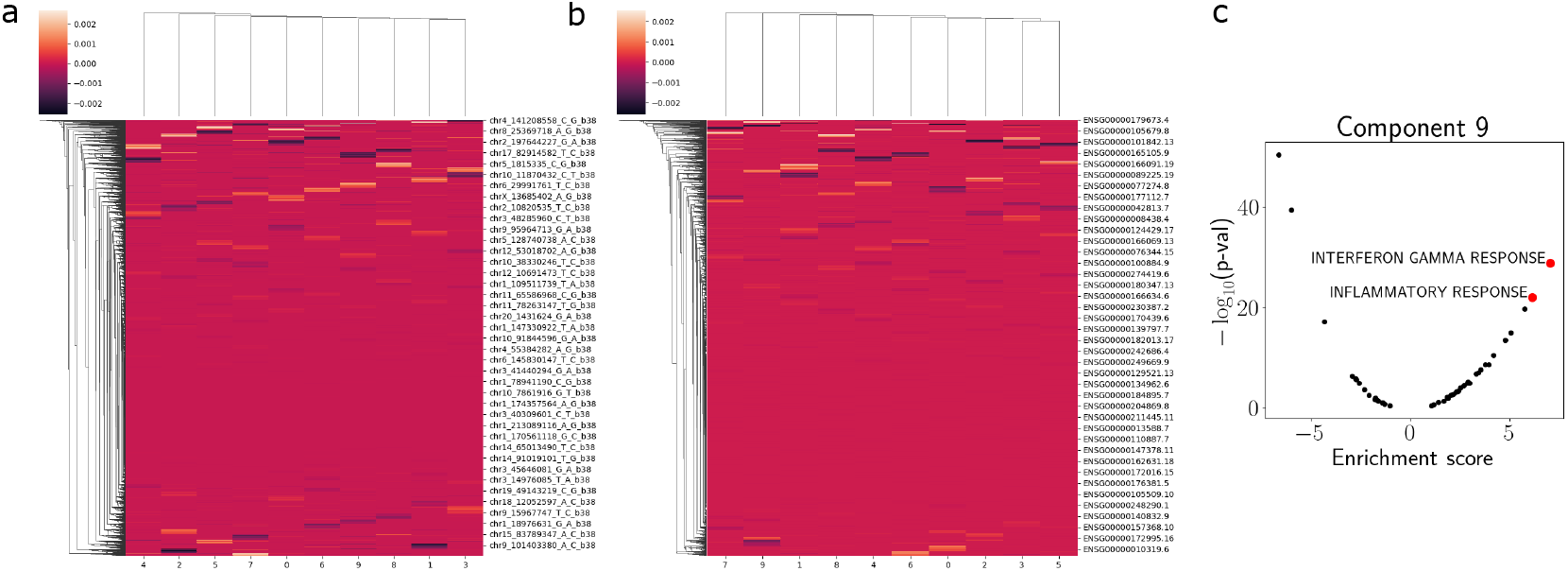
PRRR loadings matrices for the GTEx eQTL experiment. (a) A heatmap representation of the matrix **U**, showing SNPs on the rows and latent dimensions on the columns. (b) A heatmap representation of the matrix **V**, showing genes on the rows and latent dimensions on the columns. (c) Gene set enrichment analysis (GSEA) of component 9 in **V**.

**Figure 10:**
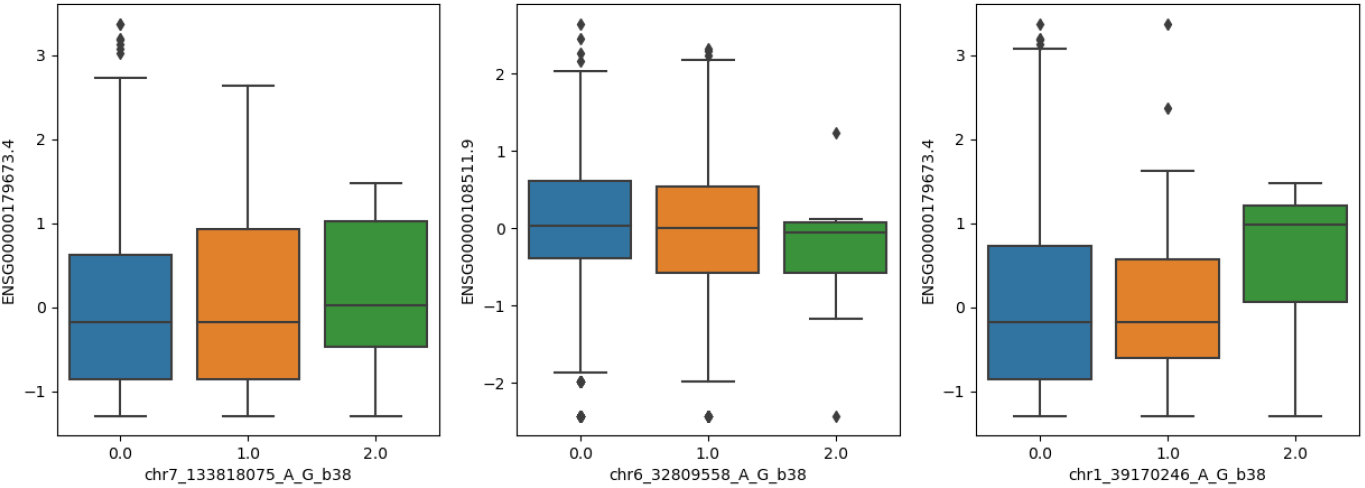
Three eQTL associations found in one latent factor of PRRR applied to GTEx liver samples.

## 4 Discussion

In this paper, we present two reduced-rank regression models and associated variational inference approaches — Poisson RRR (PRRR) and nonnegative Poisson RRR (nn-PRRR) — to jointly model associations within two high-dimensional paired sets of features where the response variables are counts. In simulations, PRRR and nn-PRRR are able to effectively capture associations between paired high-dimensional data. Moreover, we show that these models are robust to high-dimensional data, and can identify the optimal rank for the parameter matrix. In the context of sequencing data, we find that PRRR and nn-PRRR may be used for robust identification of cell types, quantifying the relationships between cell types, and performing association mapping of genetic variants to correlated genes.

Several extensions of the model could be considered. A nonparametric prior could allow for flexibly learning the rank of the parameter matrix, rather than requiring the rank to be pre-specified, as in related work (Valente et al., 2015). Additionally, the generalized model could be extended to different likelihood distributions. Furthermore, additional structure could be added to the latent variables, such as sparsity or a gene network (Engelhardt and Adams, 2014; Elyanow et al., 2020), to encode additional known structure in the covariates.

## 5 Conclusions

We present a Poisson reduced-rank regression (PRRR) model, along with a nonnegative counterpart called nn-PRRR, for association mapping in count-based sequencing data. PRRR is able to detect associations between a high-dimensional response matrix and a high-dimensional set of predictors by leveraging low-dimensional representations of the data. Using principled Bayesian modeling, PRRR is able to properly account for the count-based nature of single-cell RNA sequencing data using a Poisson likelihood. We ensure that inference is tractable and efficient in these models by applying a fast variational inference approach.

## 6 Acknowledgements

We thank Ariel Gewirtz and Isabella Grabski for helpful conversations.

## 7 Funding

This work was funded by Helmsley Trust grant AWD1006624, NIH NCI 5U2CCA233195, NIH NHLBI R01 HL133218, and NSF CAREER AWD1005627.

## 8 Code and data availability

### 8.1 Code

Code is available at https://github.com/tianafitz/PRRR.

### 8.2 Data

- Pancreas scRNA-seq data were downloaded from GEO (GSE84133). Link.
- GTEx bulk gene expression and SNP data were downloaded from the GTEx portal. Link.
- Spatial gene expression data were downloaded from the 10x Genomics “Datasets” page. Link.

## 9 Competing interests

BEE is on the SAB of Creyon Bio, Arrepath, and Freenome.

## 10 Authors’ contributions

TF, AJ, and BEE designed the method. TF implemented the method and conducted data analysis. TF, AJ, and BEE analyzed the results. TF and AJ wrote the manuscript. TF, AJ, and BEE edited the manuscript.

## 11 Appendix

### 11.1 RRR

The optimization problem for reduced-rank regression (RRR) is

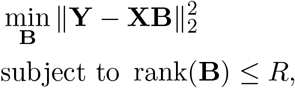

where **X** is an *N* × *P* matrix of covariates, **Y** is an *N* × *Q* matrix of outcomes, and **B** is a *P* × *Q* coefficient matrix with rank at most *R*. When *R* = min(*P, Q*), we have the OLS solution,

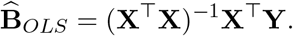

We can write the RRR optimization program in terms of the OLS solution:

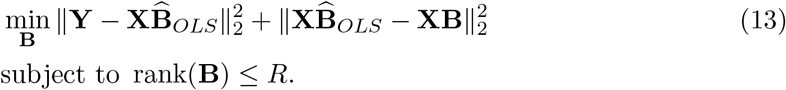

To see that these are equivalent, we can expand the loss function as follows.

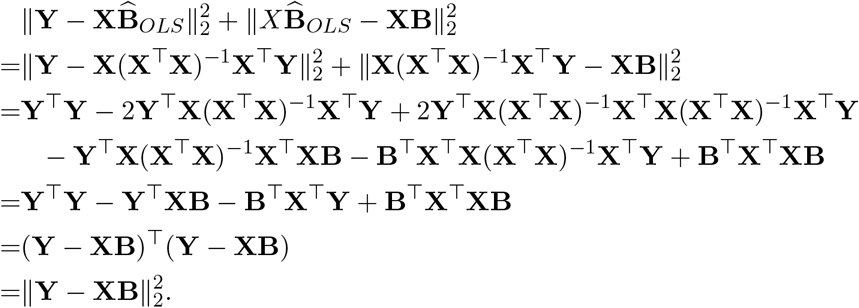

Now, the first term in the minimization problem in Equation 13 does not depend on *B*, so the problem becomes

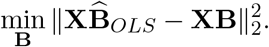

Rewriting the objective, we have

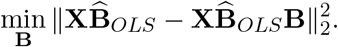

Now, we can notice that this problem aligns with PCA. Let

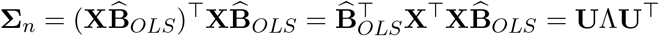

be the eigenvalue decomposition of the covariance of the fitted values. Then we have

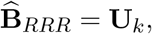

where **U**_*k*_ = [**u**_1_, · · ·, **u**_*k*_] contains the eigenvectors corresponding to the top *k* eigenvalues.

### 11.2 Plate diagrams

### 11.3 GTEx metadata experiments

In an attempt to explore the organization of PRRR’s low-dimensional space, we ran the model on a dataset containing gene expression counts along with patient metadata including height, weight, underlying conditions, and demographics. We wanted to study how features in the metadata are correlated with each other and with gene expression by analyzing the lower-dimensional matrices produced by the model. We were unable to recognize any apparent associations in the low-dimensional space, likely due to the small number of samples present in the data (about 200), and even less of these had all metadata points present.

**Figure 11:**
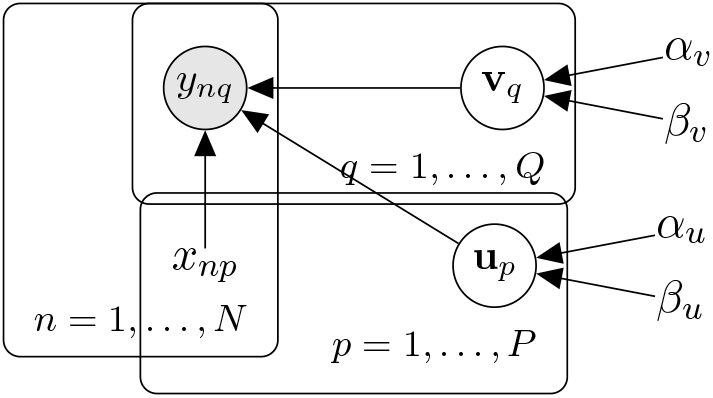
Graphical model for PRRR and nn-PRRR.

**Figure 12:**
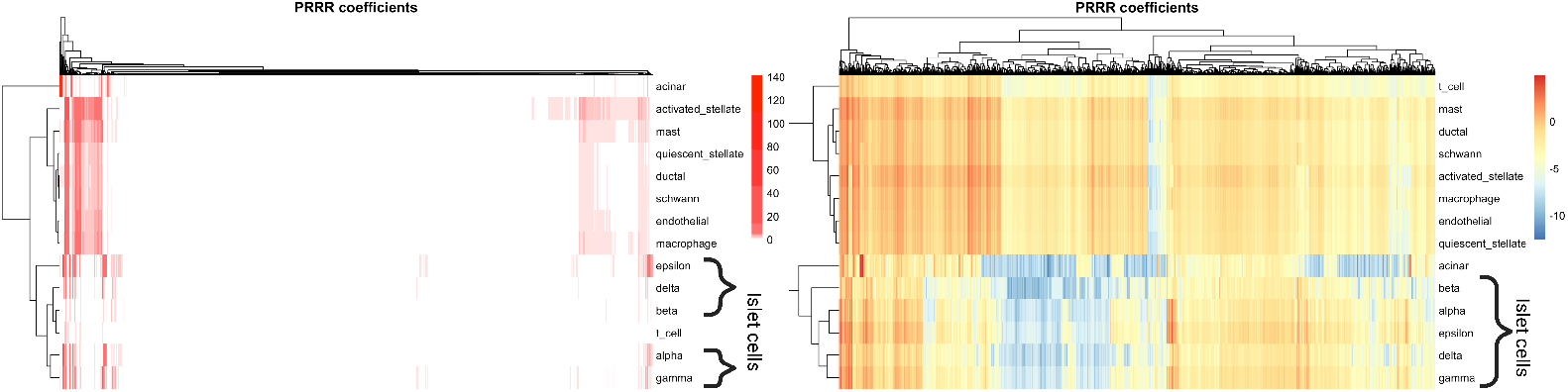
nn-PRRR coefficients for pancreatic cell types. Heatmaps showing the full coefficient matrix **UV**^*T*^ for nn-PRRR (left is original, and right is on a log scale). Cell types are shown on the rows and genes on the columns. In the left panel, white cells indicate values near zero, implying that this coefficient matrix is highly sparse.

### 11.4 Supplementary figures

**Figure 13:**
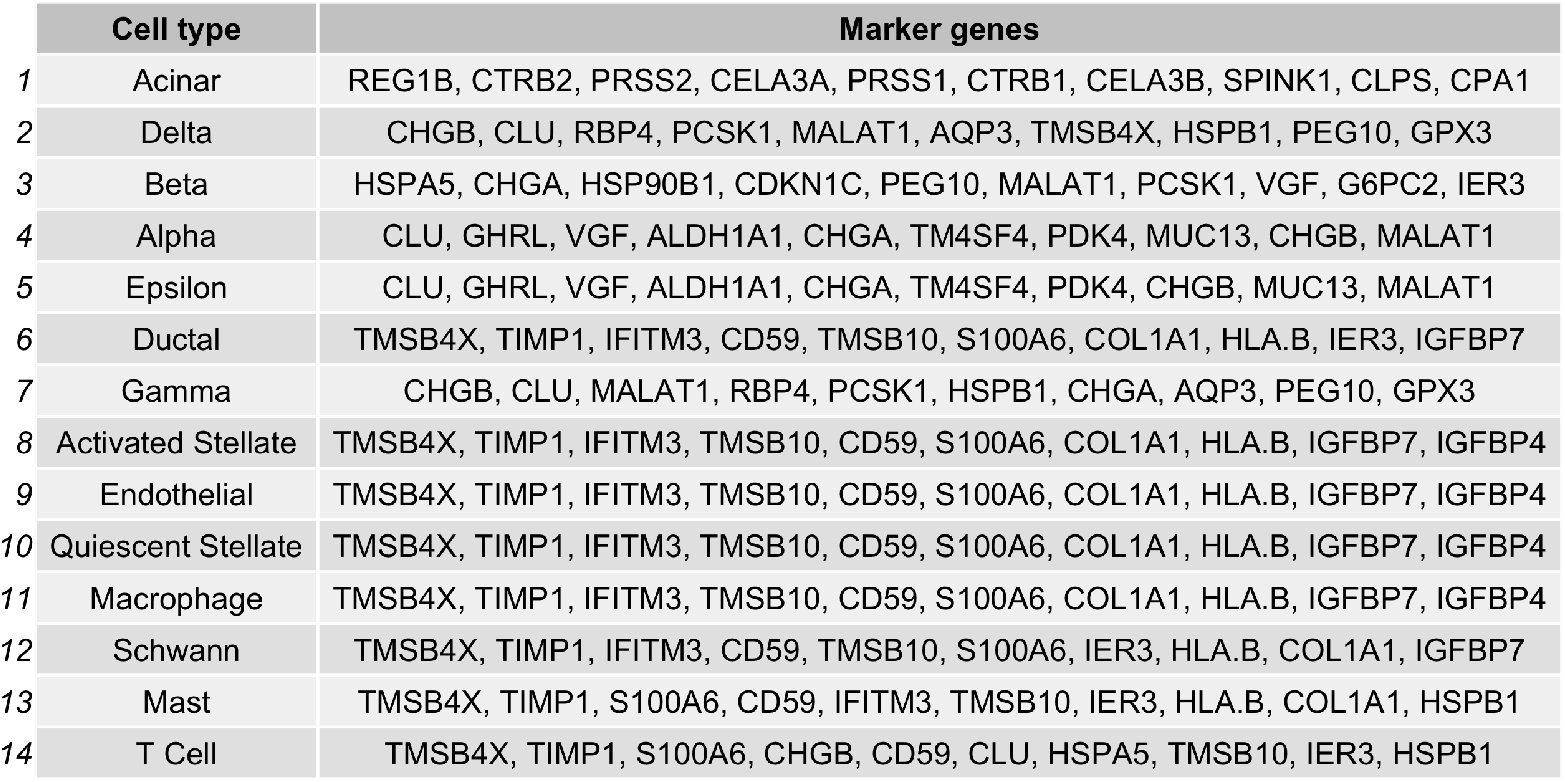
Marker genes identified by PRRR for pancreatic cell types. For each cell type, the ten genes with the highest coefficients in the matrix **UV**^T^ were extracted for each cell type. Some cell types share the same ten marker genes, which corresponds with our observation that the cell types are largely overlapping in a PCA plot of the gene expression data (Figure 14).

**Figure 14:**
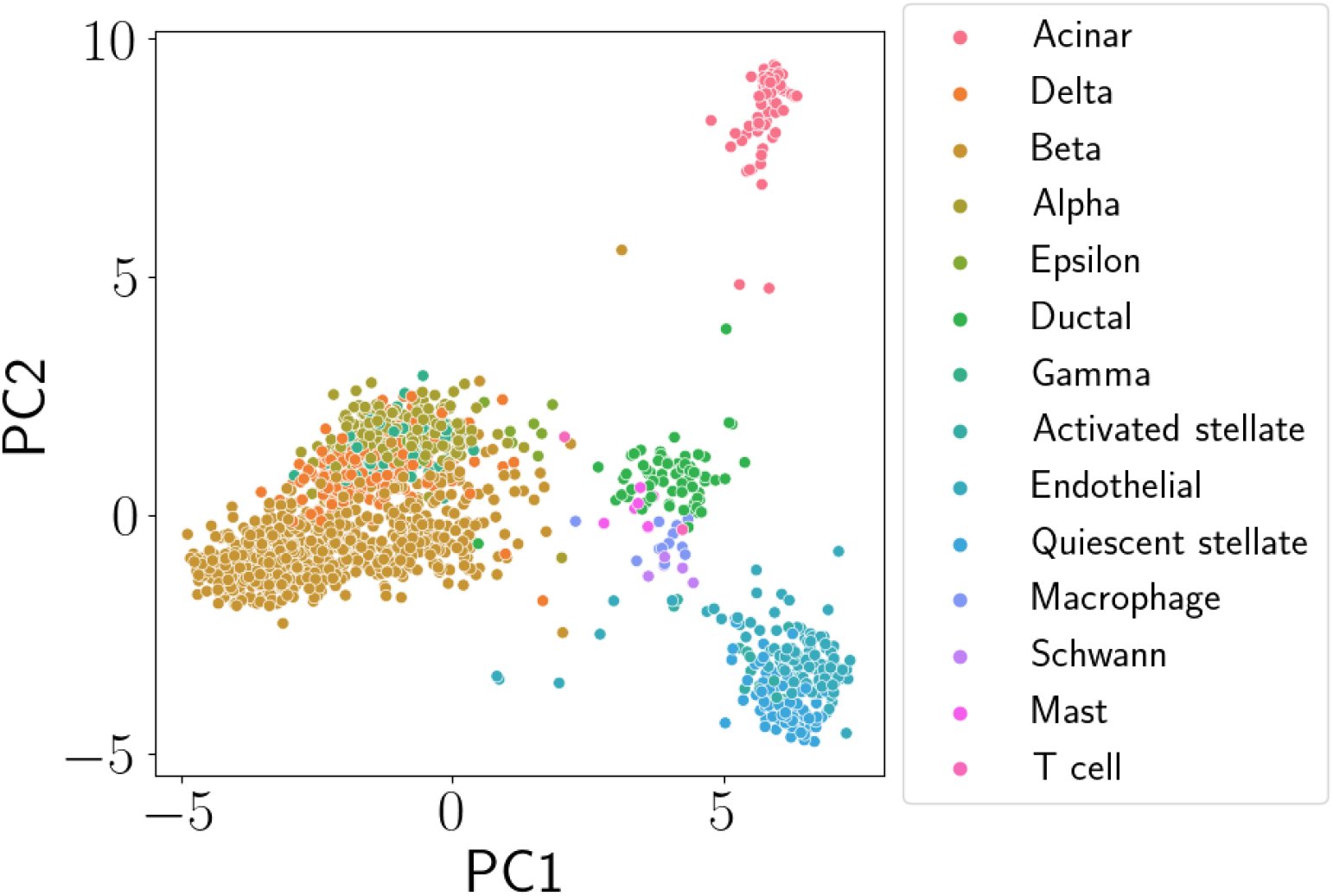
PCA plot of pancreas scRNA-seq data. The first two principal components (PCs) are plotted. Each point corresponds to a single cell and is colored by its annotated cell type.

**Figure 15:**
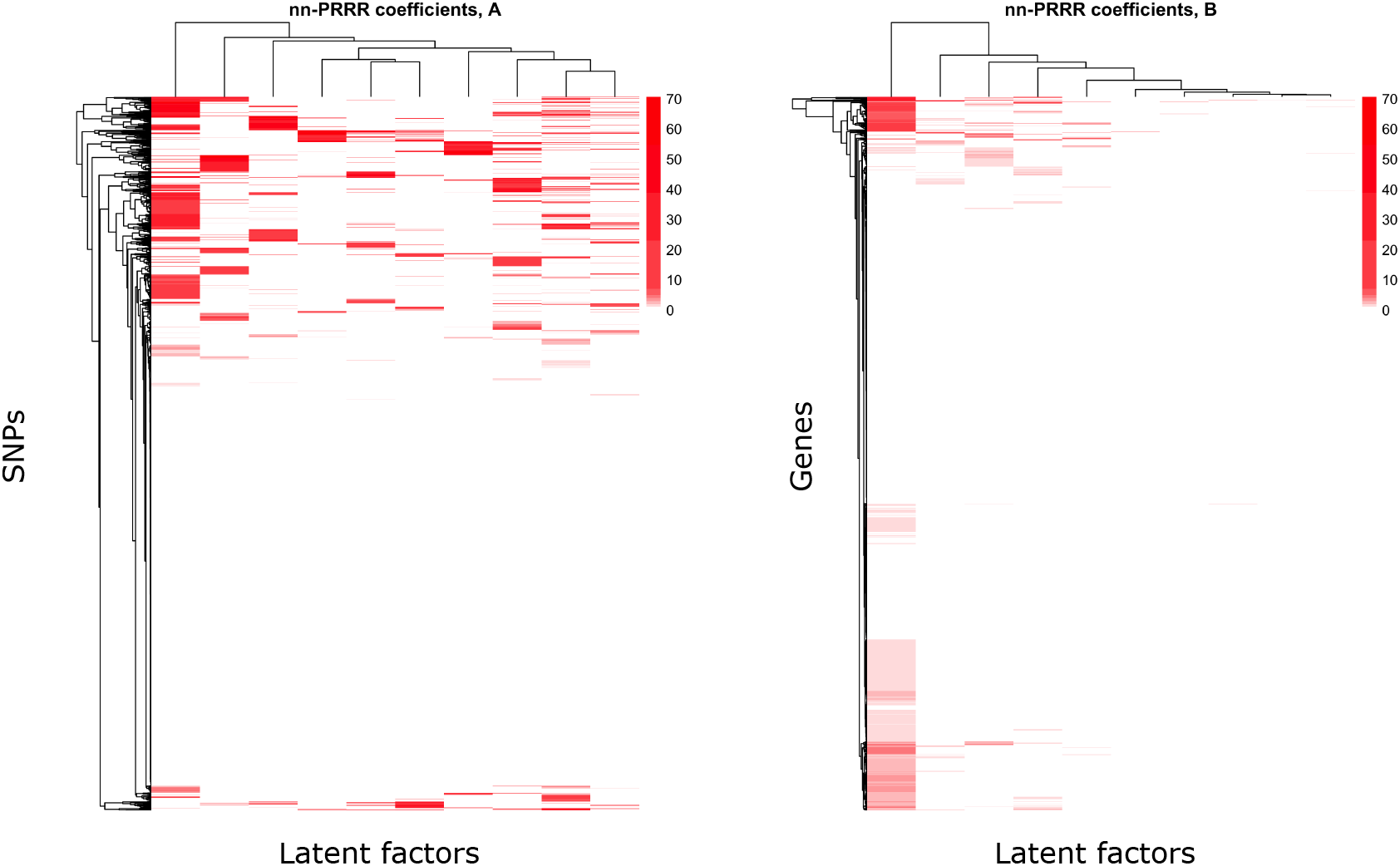
nn-PRRR coefficients for GTEx eQTL mapping. Left: **U** matrix showing SNPs on the rows and latent factors on the columns. Right: **V** matrix showing genes on the rows and latent factors on the columns.

**Table 1:**
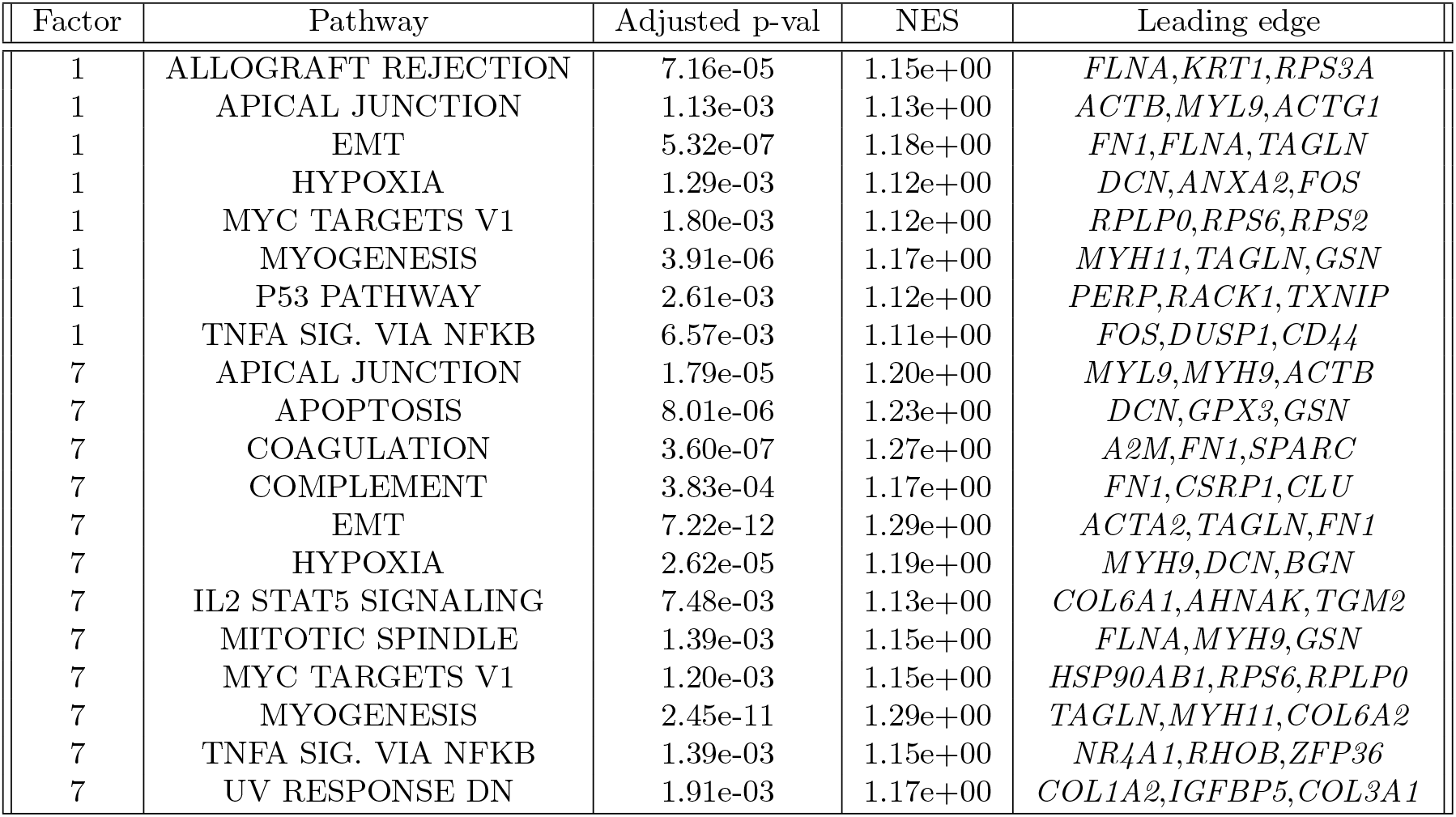
Gene set enrichment results for GTEx eQTL experiment. *EMT* stands for epithelial mesenchymal transition.

